# Phenotypic determinism and contingency in the evolution of hypothetical tree-like organisms

**DOI:** 10.1101/527622

**Authors:** Tomonobu Nonoyama, Satoshi Chiba

## Abstract

Whether evolutionary history is mostly contingent or deterministic has been given much focus in the field of evolutionary biology. Studies addressing this issue have been conducted theoretically, based on models, and experimentally, based on microcosms. It has been argued that the shape of the adaptive landscape and mutation rate are major determinants of replicated phenotypic evolution. In the present study, to incorporate the effects of phenotypic plasticity, we constructed a model using tree-like organisms. In this model, the basic rules used to develop trees are genetically determined, but tree shape (described by the number and aspect ratio of the branches) is determined by both genetic components and plasticity. The results of the simulation show that the tree shapes become more deterministic under higher mutation rates. However, the tree shape became most contingent and diverse at the lower mutation rate. In this situation, the variances of the genetically determinant characters were low, but the variance of the tree shape is rather high, suggesting that phenotypic plasticity results in this contingency and diversity of tree shape. The present findings suggest that plasticity cannot be ignored as a factor that increases contingency and diversity of evolutionary outcomes.

## 1. Introduction

There has been much debate as to whether the evolutionary history of life is mostly contingent or more or less deterministic [1–4]. A metaphor of “replaying life’s tape” [1] was used to emphasize the preeminent role of contingency in the evolutionary process [1]. In this view, the outcome of evolution could be dramatically different from the actually observed course of events, because evolution is essentially a stochastic phenomenon whereby trajectories that start infinitely close to each other soon diverge (because divergence is exponential). Experimental study of bacteria has suggested that small coincidences of history might lead populations along different evolutionary paths [5]. On the other hand, it was claimed that natural selection constrains organisms to a relatively few highly adaptive options [3]. In this view, the evolutionary routes are many, but the destinations are limited. In the present study, we examine relative contributions of determinism and stochasticity in evolution. This has not been addressed directly, and the quantification of the predictability of evolution remains elusive.

The concept of a fitness landscape [6] has been used to address this issue. This concept has influenced many research fields on evolution and much effort has been spent to understand the characteristics of empirical landscapes.[7–9]. The base concept of a fitness landscape assumes that there is a functional relationship between the genome of an organism and its growth rate and fitness [10]. The model of the fitness landscape describes how some phenotypes are more likely to evolve than others, and how developmental mechanisms could limit the evolutional change [11]. In this situation, multiple evolutionary trajectories are accessible, but evolution might be strongly constrained to a particular adaptive peak [12].

However, in many real-world scenarios, fitness evaluations are not trivial [13] and experimental evolution involving sexual reproduction and multicellular organisms with long life timescale is difficult [14], so that sometimes it is difficult to apply the fitness landscape into real biological systems. Also, phenotypic changes are often largely affected by phenotypic plasticity. The shape of the adaptive landscape might largely be affected by phenotypic plasticity, while such effects of plasticity have not been considered in the studies of repeatability of phenotypic evolution. In the present study, we simulate parallel evolution experiments that focus on the predictability of evolution while also considering the effects of plasticity.

To address this subject, a community of dendritic organisms (e.g., tree, coral, sponge, Stromatoporoidea) provides an excellent model [15–18]. We assume hypothetical tree-like organisms and simulate their evolution and plastic change using the Honda model [19]. Using this model, we construct an individual-based model in which individuals compete with each other for light. We assume organisms are growing under direct ambient light, and we are not strictly concerned about the weight of their body. The morphology of the organisms was assumed to be determined by plastic change as a response to environmental condition and by genetic change as a result of genetic drift, mutation, and natural selection. For example, the morphology of coral is determined by both environmental factors, particularly light environment [20,21] and genetic components [22]. We assume that the common evolutionary patterns can be found by simulating the shape of the hypothetical tree-like organisms [23].

This study aims to explore the factors that affect the evolutionary contingency and determinism. We test whether, after sufficient time, the same phenotypes evolve in the model community within every simulation. Particularly, we focus on how mutation rates affect the contingency of the evolutionary history and how plasticity is associated with the phenotypic diversity patterns.

## 2. Methods

Here we constructed an individual-based model, in which dendritic-shaped individuals compete for light with other individuals and evolve through mutation and natural selection. Each individual was described by simple branching rules and parameters (e.g., Honda 1971). Using these models, the optimal branching rule to minimize the overlapping of leaf clusters was obtained [24,25]. For convenience, the organism in the model was called a “tree” and its single component was called a “branch”. In the present tree model, we tentatively call the clusters of trees a “forest”.

In this model, the branch development process was deterministic. The variety of tree shapes was produced by the growth of branches to adapt to the surrounding light environments. To describe how leaves receive light and how it inhibits other leaves from receiving light, a leaf ball model was used [26]. This model considers a leaf cluster as a ball and the branch according to the brightest vector. The influence of the local light environment on branch growth is an essential process determining the dynamics of tree crown architecture [27,28]. Using these methods, we developed the simulation model, incorporating the dependence of the number and size of new shoots on the photosynthetic production of parental shoots. This model describes the growth of trees by branch developments, which is determined by the light condition. Accordingly, in this study, we used a previous 3D tree growth model [19,29] by incorporating a leaf ball model and reaction process of branch developments against the photosynthetic condition.

### 2.1 Tree growth

The tree morphology is determined by 11 parameters (Table 1). These parameters are all used by Honda[19] and Yoshizawa [30] and the basic correlation between the tree shape and parameters are already shown so that that will not be exactly analyzed in this paper. The exact description of the branching geometry is as follows. A parent branch diverges into two offspring branches through one branching process. The length of each of the two offspring branches is determined by the ratio of *R*_1_ and *R*_2_, respectively, to the length of the parent branch. The angle between the mother branch and two offspring branches are described as *θ*_1_ and *θ*_2_ respectively. In the case of the trunk, when the *θ*_1_ or *θ*_2_ equal to zero, one of the offspring branches grows vertically. Another branch was assumed to form a divergence angle alpha with the sister branch of its parent branch (i.e., the offspring branch is rotated with angle alpha around the trunk from the sister branch of its parent branch).

**Table 1.** Parameters used in the simulation.

| Parameter | Description | Range |
| --- | --- | --- |
| $\theta_1$ | Branching angle 1 | $0 \leq \theta_1 \leq \pi$ |
| $\theta_2$ | Branching angle 2 | $0 \leq \theta_2 \leq \pi$ |
| $br_1$ | Branch attenuation rate 1 | $0 < br_1$ |
| $br_2$ | Branch attenuation rate 2 | $0 < br_2$ |
| $\alpha$ | Rotation angle | $0 \leq \alpha \leq \pi$ |
| $lr$ | Leaf radius | $0.0 < lr$ |
| $la$ | Leaf absorption rate | $la < 1.0$ |
| $cr$ | Tendency to move | $-1.0 < cr < 2.0$ |
| $at$ | Avoid threshold | $0.0 < at < 1.0$ |
| $ra$ | Rate of avoid reaction | $0.0 \leq ra \leq 1.0$ |
| $dt$ | Death threshold | $0.0 < dt < 1.0$ |

It is assumed that the branches search for the direction of light in the surrounding environment. To describe a cluster of leaves, this model places a ball at the distal end of each branch. This ball is used to determine the brightest direction by making a map of the shadow of the other branches on the ball surface. The direction in which the branches will grow was determined according to the incoming light vector. ***V*** is the average vector of light. ***V*** is calculated by the following equation (Kanemaru et al, 1992): ***V*** = (Σ_*n*_Σ_*m*_*l*(*n,m*)***N***(*n,m*)*S*(*n,m*))/|Σ_*n*_Σ_*m*_*l*(*n,m*)***N***(*n,m*)*S*(*n,m*)| where *l*(*n,m*) is the brightness per area, ***N***(*n,m*) is the vector from the center of the ball to each cell on the ball surface, and *S*(*n,m*) is the area on the ball surface. n and m describe longitude and latitude. The light amount (*L*_*e*_) received by the leaf is calculated by the following equation: *L*_*e*_ = Σ _*n*_Σ _*m*_*l*(*n,m*)*S*(*n,m*). Once the amount of the light received by the tree is calculated, branches adjusted their positions according to ***V***. The considered branch is rotated around the point where it attaches to its parent branch, toward brightest direction. The actual angle used for changing branch direction, *Ø*, is calculated by *Ø* = *crδ*, where *δ* is the angle between ***V*** and original direction of the branch to grow. *cr* is the constant rate of bending the branch. In the case of *cr* = 0.0, there is no influence of the phototropism in the simulation. After all branches change their direction, those branches that cannot capture sufficient light do not extend and become lost. Even if the branches can capture sufficient light to grow, the growth rate of the sister branch changes with the ratio of *ra* [30]. The amount of light captured by an individual tree is calculated by summing all light captured by the survived branches of the individual tree. The light amount captured by the *Nth* tree is calculated by the equation *L*_*N*_ = Σ*L*_*e*_. Trees are placed in a square area at an even interval. The ground of the area is assumed to reflect light at a fixed rate. Light is considered as ambient light.

### 2.2 Reproduction and evolutionary process

The simulation area is divided into the number of square cells. Each tree grows within a cell of the simulation area. The trees produce seeds and have the same lifespan. There is no overlap of generations among individuals. In the model, we assume that one tree produces one seed (offspring), and individuals with low fitness cannot produce seed (offspring). The total number of individuals is assumed to be constant.

We assume that the trees are all clones, and the 11 variables determining tree morphology are the same between offspring and its mother, except in cases of mutation, which occurs with a defined probability. A change of the value of each variable by mutation is described by adding a random value having a Poisson distribution with a mean of 0 and a standard deviation of 1.

After each generation, all of the cells are assumed to become empty. Each cell is placed by an offspring following the order of fitness of the offspring produced by all individuals. If the fitness is equal, the offspring is selected at random and placed on the cell. The fitness *F* of the *Nth* individual is calculated using the following equation: *F*_*N*_ = *L*_*N*_ *X*_*N*_(K - *X*_*N*_)*A*_*N*_(P - *A*_*N*_) where K and P are the constant, *L*_*N*_ is the light amount captured by *Nth* individual, *X*_*N*_ is the whole length of the individual, and *A*_*N*_ is the value calculated by multiplication of the size of the tree and efficiency of the light. Durable construction and efficiency cannot be achieved at the same time [31]. In this paper, we consider the durability of leaves to be correlated with the size of leaves; because it seems that a large leave needs a structure to support it.

### 2.3 Phenotypic difference

We evaluated the tree phenotypes using two parameters, the aspect ratio of the branches and the number of branches of each individual. These two parameters are considered to be independent values. These parameters were calculated for all individuals, and their variance and average were obtained. Also, we examined changes in 11 genetically-determined variables, and calculated the average and variance for each variable. After sufficient generation times, the phenotypes of the evolved trees were compared among the different simulation runs, and thus, we can investigate if similar phenotypes evolve independently in the different simulation runs. At the end of each simulation run, the values of the above two parameters were plotted in the two-dimensional space. We compare the patterns and position of the plots among different simulation runs. We assume overall phenotypic difference *D*. The overall phenotypic difference between the *Ath* simulation run and the *Bth* simulation run are calculated as 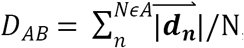, where ***d***_*n*_ is the shortest vector from *nth* plot in the *Ath* run to the *mth* plot in the *Bth* run. The average of the shortest distance of this vector represents the overall phenotypic difference between the different simulation runs.

## 3. Results

We examined the aspect ratio and number of the branches of the all trees generated after 2500 generations (Fig. 1, Fig. 2). The number of types with different values for the aspect ratio of the branches and number of branches was larger with the higher mutation rate, and every individual has unique values for these traits at *μ* = 5.0*e* - 1. Due to plasticity, there is no fixed form tree. The shape of the individuals is highly influenced by their surrounding environment. The relationship between mutation rate and average minimum distance of the phenotypic distribution among the different simulation runs is shown in Figure 3. The differences in the phenotypic distribution among the different simulations represented a hump-shaped relationship with the mutation rate, but its peak is located near the lowest mutation rate (5 × 10^−5^). The difference became the lowest at the highest mutation rate. Both of the variances of the aspect ratio of the branches and number of the branches became the lowest at the intermediate level of mutation rate (Fig. 4). In contrast, the variance of the 11 characters showed a hump-shaped or monotonical increase with increasing mutation rate (Fig. 5). The differences among these patterns of variance reflect the differences in the variances of phenotypic plasticity. Thus, the variance of phenotypic plasticity became rather high at low mutation rates (5× 10^−5^ – 5 × 10^−6^). The shape of the tree was the most complex at the 5 × 10^−5^ mutation rate, and became rather simple at the high mutation rate.

**Fig. 1.**
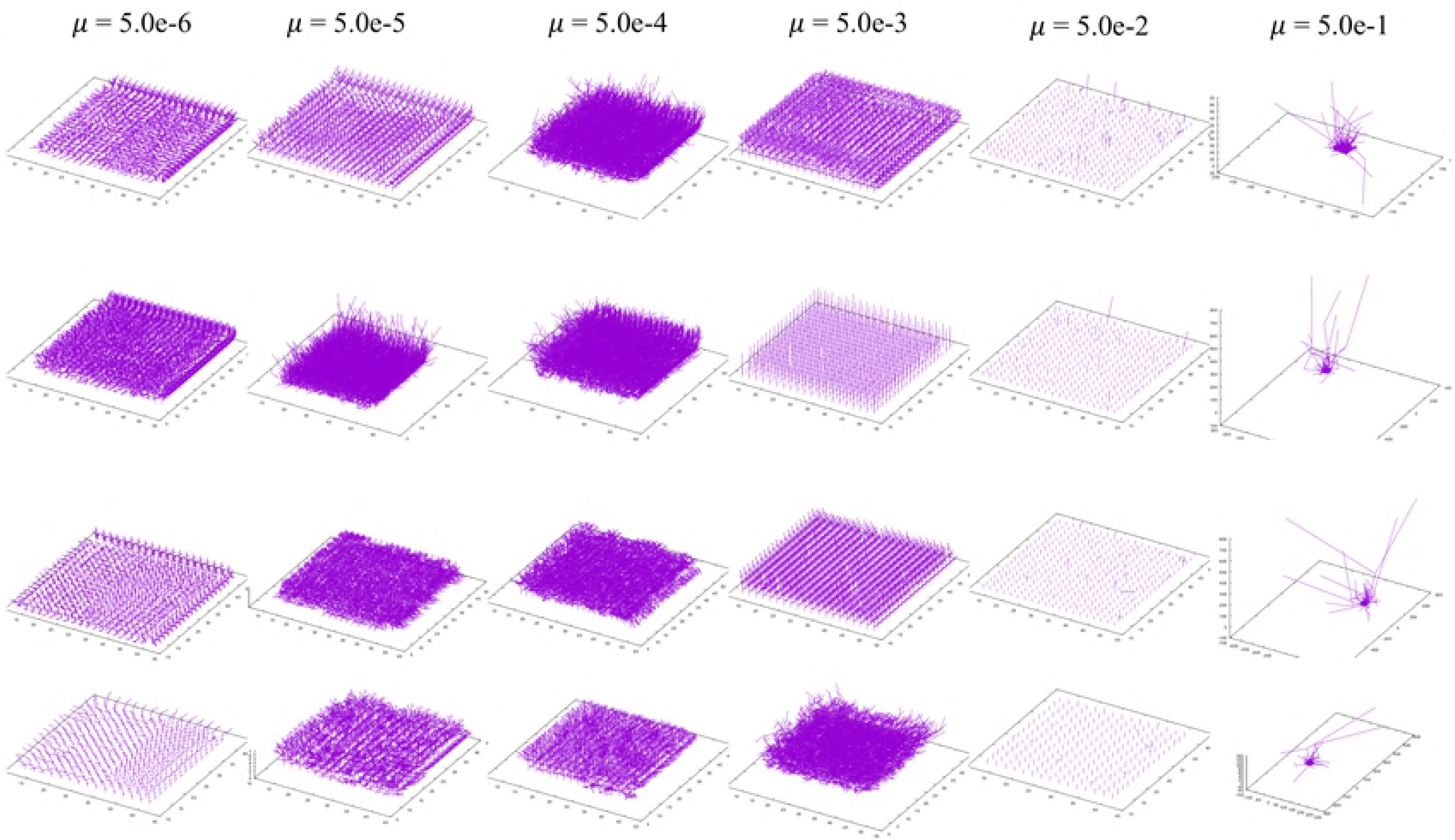
The 3-D shape of the results. Each map is one experiment result. The six rows represent the mutation rates. The mutation rates were 5.0e-1,5.0e-2,5.0e-3,5.0e-4,5.0e-5,5.0e-6, from the left. The three columns represents the individual number. The bottom column is the result which contains 225 individual and other columns is in the result containing 400 individuals. All results are after 2500 time steps.

**Fig. 2.**
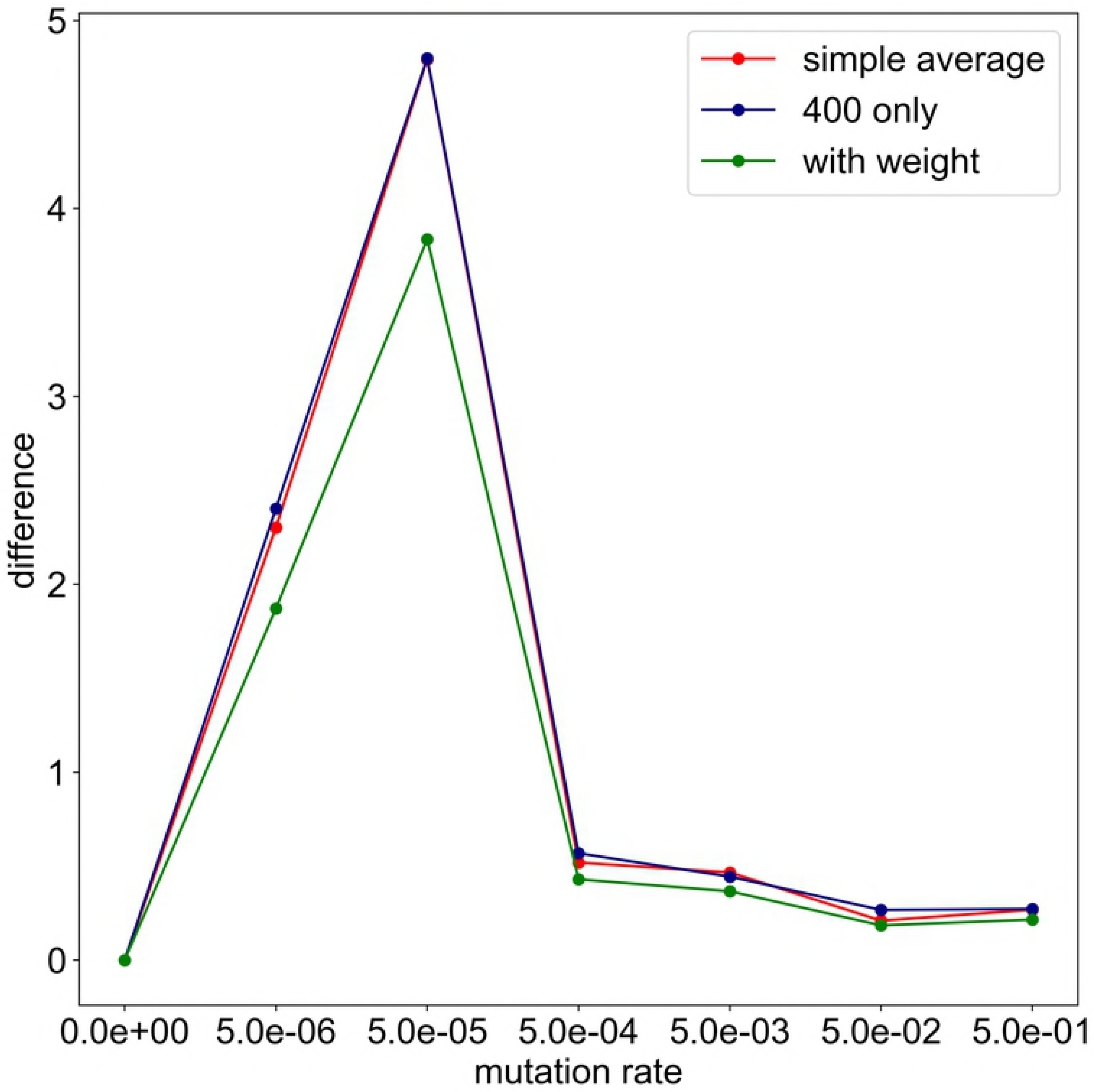
Architecture of model organism after 2500 generations. Each individuals make branch for 5 times. The mutation rates were 5.0*e* - 1, 5.0*e* - 2, 5.0*e* - 3, 5.0e - 4, 5.0e - 5, 5.0e - 6, from the left. The bottom column is the result which contains 225 (15 × 15) individual and other columns is in the result containing 400 (20× 20) individuals.

**Fig. 3.**
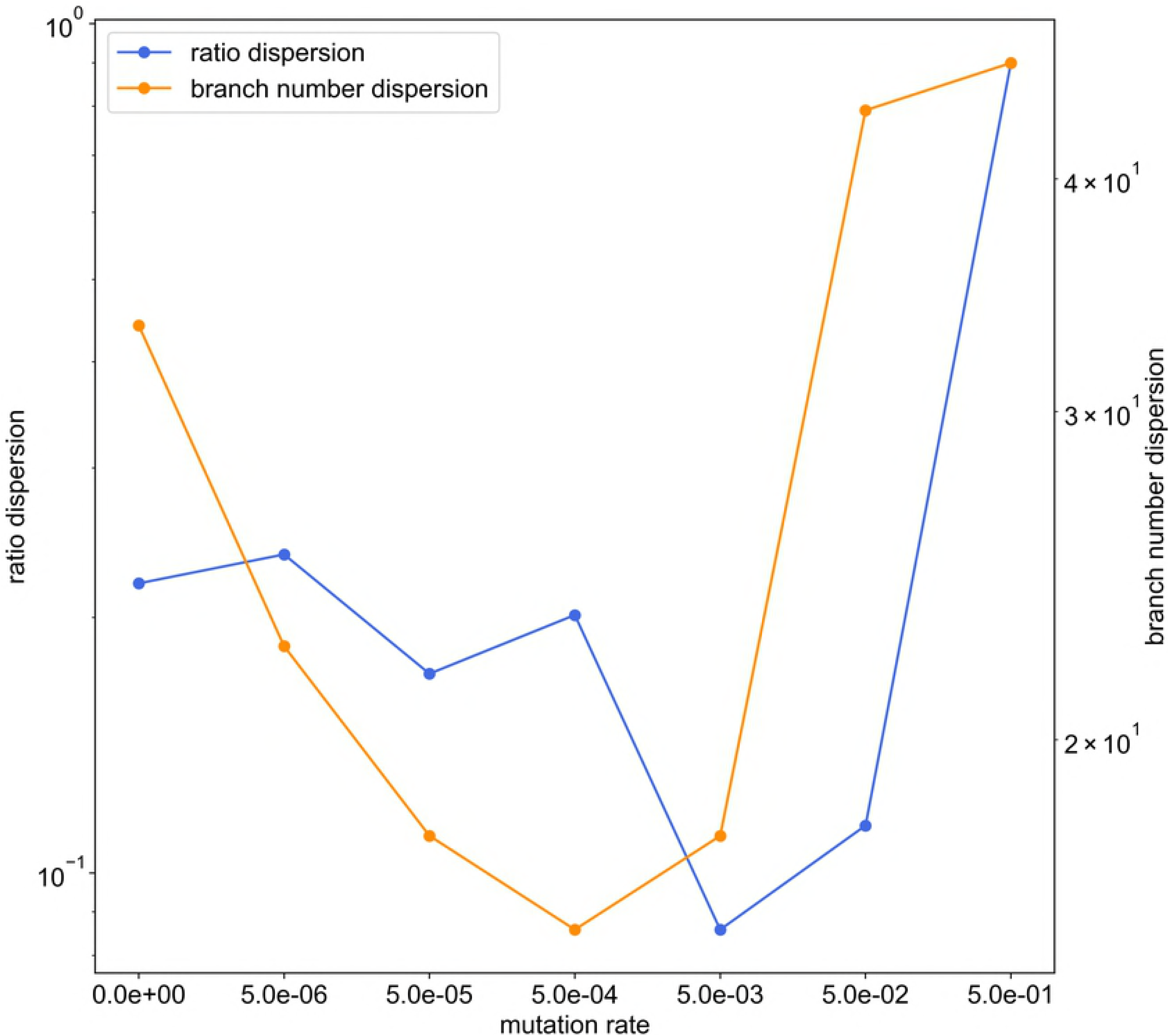
The histogram of the whole result. Each map is one experiment result, and the horizontal axis represents aspect ratio, which is calculated by *D*_*v*_/*D*_*h*_, where *D* is the distance from the root to the most furthest branch, and small v and h denote the horizontal distance and vertical distance. The vertical axis represents the number of the branches in the individual. The six rows represent the mutation rates. The mutation rates were 5.0*e* - 1, 5.0*e* - 2, 5.0*e* - 3, 5.0*e* - 4, 5.0*e* - 5, 5.0*e* - 6, from the left. The three columns represents the individual number. The bottom column is the result which contains 225 individual and other columns is in the result containing 400 individuals. All results are after 2500 time steps. The color bar of the figures are limit at 10 individuals.

**Fig. 4.**
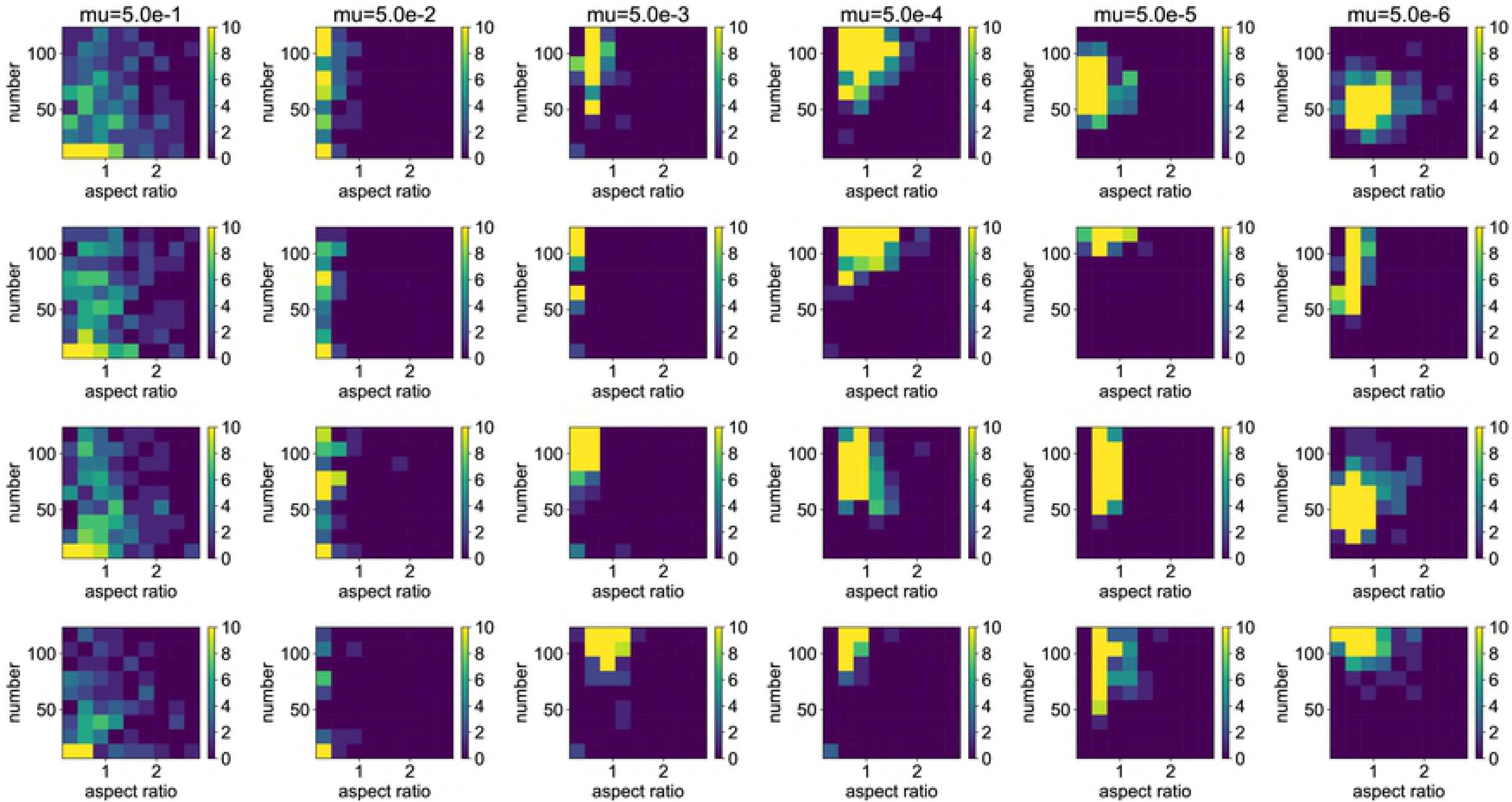
The average of the minimum distance. The horizontal axis represents the mutation rate. The vertical axis represents the average of the minimum distance cases of the same mutation rate. The green line represents for the simple average of four cases, and the blue line represents for the average which is considering the number of the individuals and weight. The orange line represents for the result only in 400 individuals. If the simulations all end at the same result, all points in the histogram become the same, and the minimum distance equals zero because all points are overlapped.

**Fig 5.**
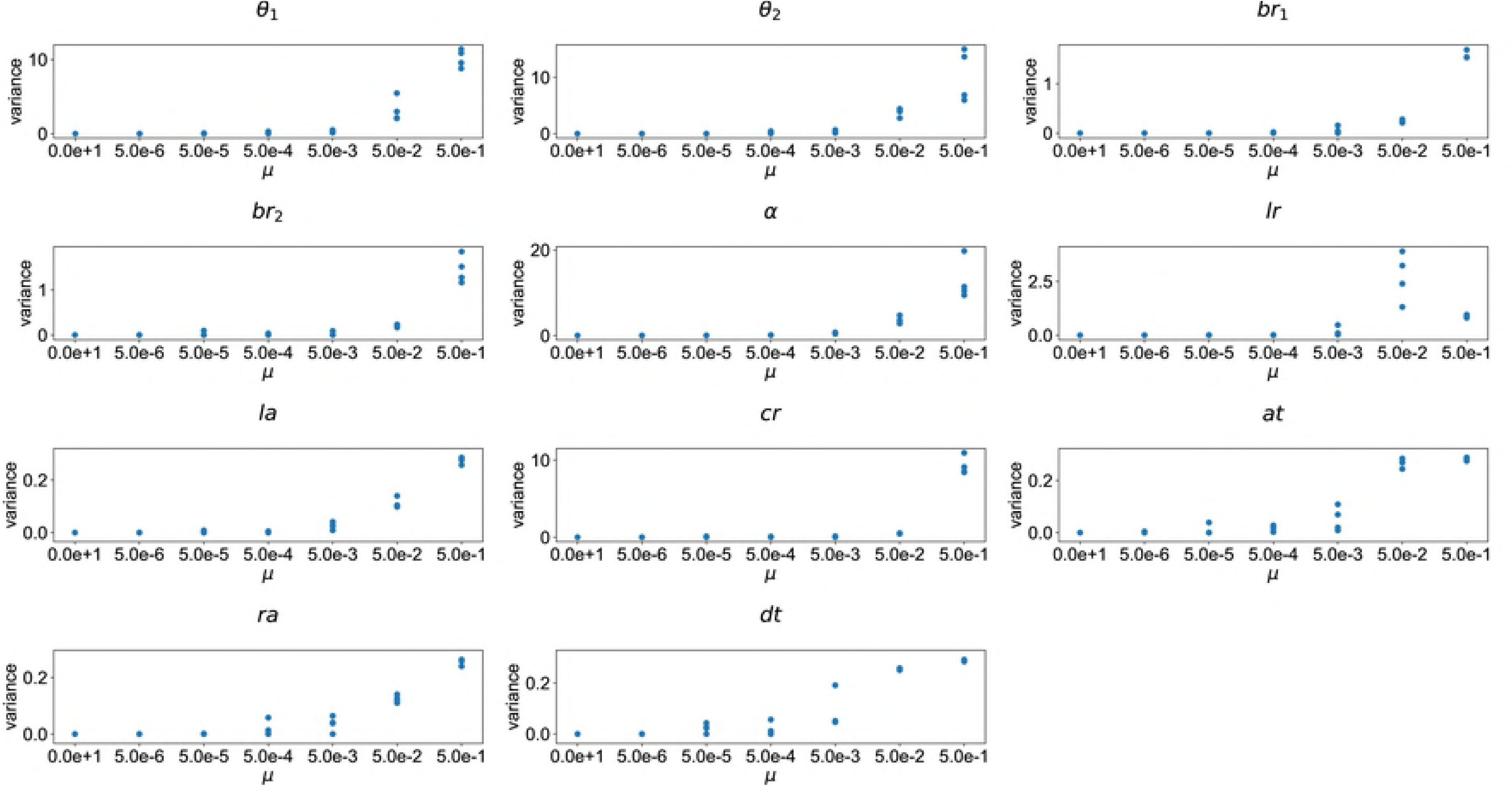
The Variance of the result in aspect ratio and branch number. The horizontal axis represents the mutation rate. The left vertical axis represents the aspect ratio dispersion, and the right vertical axis represents the branch number dispersion. Both lines are a downward convex.

Fig 6. The variance of the parameters.

The parameter meanings are shown in Table 1. The horizontal axis represents the mutation rate. The vertical axis represents the variance of each parameter.

## 4. Discussion

A high mutation rate enables the phenotype to more easily move on the surface of the adaptive landscape. Under the high mutation rate, the population consists of multiple competing clones that differ genetically from each other [32,33]. In these cases, the phenotype reaches a global peak in the adaptive landscape [12]. Therefore, it is expected that the same phenotype evolves at the end of all simulations when the mutation rate is high. In terms of the two phenotypic characters, including both effects of genetic components and plasticity (aspect ratio of branches and number of branches), the results of the present simulation are consistent with this expectation. However, in this simulation, the variances of these traits are the highest at the highest mutation rate. The variances of many of the genetically determined characters also became highest at the highest mutation rate. These findings mean that phenotypic characteristics are mostly determined by the genetic component when the mutation rate is very high. A high mutation rate (e.g., 5 × 10^−3^ and 5 × 10^−4^) causes a decrease of branch number and aspect ratio, resulting in thin and sparse individuals. These characteristics are the least susceptible to plastic change. The number of branches is higher at the mutation rate of 5 × 10^−2^ than at lower mutation rate. Quite small and many branches were developed in several trees under the condition of this mutation rate, so that aspect ratio has not much changed relative to the lower mutation rates. Because small branches hardly interfere with the other branches, this type of tree also does not represent plastic change. In this case, the mean phenotype evolved is mostly equal among the different simulations, but the variation of the phenotype is high due to the high mutation rate.

In contrast, a high variety of phenotypic distribution patterns arises at the mutation rate of 5 × 10^−5^. A low mutation rate implies that it is difficult to move a long distance on the fitness landscape [34]. The dominant phenotypes of the two traits (aspect ratio of branches and number of branches) that appeared in each simulation differ among the simulations. This pattern implies that phenotypes are trapped at particular adaptive peaks, and cannot reach a global peak of the adaptive landscape. However, in this simulation, the variances of these traits, particularly of the aspect ratio of branches, becomes rather high at the mutation rate of 5 × 10^−5^. Also, variances are low in most of the genetically determined characters at this mutation rate. Thus, the phenotypic variances arising at this level of mutation rate are mainly attributed to phenotypic plasticity. Yokozawa demonstrated that crown architecture traits are important for the pattern of species coexistence in trees [35]. Complex tree shape and high variability in tree morphology are maintained due to the plasticity.

In this simulation, phenotypic plasticity results from the interactions among branches attempting to obtain light. Such interactions among branches might yield the unique shape of the tree by chance. When the mutation rate is 5 × 10^−5^, the shape of the tree becomes complex. This suggests that the interactions among the branches are serious, resulting in the complex shape of the tree. Under such conditions, small spatial differences in the light environment appear by chance as a result of interaction among branches of the same and different trees. These differences might change the shape of the adaptive landscape. Thus, such plasticity is likely to promote the evolution of divergent phenotypes. Also, plasticity increases contingency, resulting in less repeatability of the evolutionary outcome among the different simulations.

When the mutation rate is very low (i.e., 5 × 10^−6^), the repeatability of the phenotypic distribution remains low. This low level of repeatability and high level of contingency are supposedly derived from the effect of phenotypic plasticity. This is because variances of the genetically determined characters are very low and near zero, but variances of the two phenotypic traits, which include variance due to plasticity, are relatively high. Thus, phenotypic plasticity again increases the contingency and yield different phenotypic distributions at the end of the different simulations.

In the present simulation, we could not separate the phenotypic variance into genetic variance and environmental variance. We estimated the effect of the plasticity on the phenotypic variance by comparing the patterns of relationship between variables to develop the tree and parameters describing the tree shape. Although the former is determined only genetically and reflects genetic variance, the latter includes both genetic and environmental variance. Therefore, to better estimate the effects of plasticity in future studies, it is necessary to develop a model that can separate genetic variance and phenotypic variance. Furthermore, the regeneration niche has been regarded as an important factor for coexisting of species [36,37]. The overlap of individual rearrangement and generation change might also affect the coexistence of species. However, the present findings still suggest that phenotypic plasticity cannot be ignored as a factor to enhance the diversity of phenotypes through the increase of contingency. Plasticity potentially plays a major role in producing phenotypic diversity with relatively low mutation rates.

## Acknowledgements

We thank T. Aota and T. Suzuki for technical support and suggestive advice on this study. This study was funded in part by JSPS KAKENHI Grant Number 17H04611.

## References

1. Gould S. Wonderful life: the Burgess Shale and the nature of history. Trends in Ecology & Evolution. W. W. Norton and Company; 1990.

2. Vermeij GJ. Historical contingency and the purported uniqueness of evolutionary innovations. Proc Natl Acad Sci. 2006;103: 1804–1809. doi:10.1073/pnas.0508724103

3. Morris S. Life’s solution: inevitable humans in a lonely universe. Cambridge University Press; 2003.

4. Powell R. Convergent evolution and the limits of natural selection. Eur J Philos Sci. Springer Netherlands; 2012;2: 355–373. doi:10.1007/s13194-012-0047-9

5. Blount ZD, Borland CZ, Lenski RE. Historical contingency and the evolution of a key innovation in an experimental population of Escherichia coli. Proc Natl Acad Sci. 2008;105: 7899–7906. doi:10.1073/pnas.0803151105

6. Wright S. The roles of mutation, inbreeding, crossbreeding and selection in evolution, Proceedings of the Sixth International Congress of Genetics. Proc Sixth Int Congr Genet. 1932;1: 356–366.

7. Weinberger ED, Kauffman SA. The NK Model of rugged fitness landscapes and its application to maturation of the immune response. J Theor Biol. 1989;141: 211–245. Available: https://www.sciencedirect.com/science/article/pii/S0022519389800190

8. Dawid A, Kiviet DJ, Kogenaru M, de Vos M, Tans SJ. Multiple peaks and reciprocal sign epistasis in an empirically determined genotype-phenotype landscape. Chaos An Interdiscip J Nonlinear Sci. 2010;20: 026105. doi:10.1063/1.3453602

9. Lobkovsky AE, Wolf YI, Koonin E V. Predictability of Evolutionary Trajectories in Fitness Landscapes. Shakhnovich EI, editor. PLoS Comput Biol. 2011;7: e1002302. doi:10.1371/journal.pcbi.1002302

10. Gavrilets S. Fitness landscapes and the origin of species. Princeton: Princeton Univ. Press; 2004.

11. Beldade P, Koops K, Brakefield PM. Developmental constraints versus flexibility in morphological evolution. Nature. 2002;416: 844–847. doi:10.1038/416844a

12. Lobkovsky AE, Koonin E V. Replaying the Tape of Life: Quantification of the Predictability of Evolution. Front Genet. 2012;3. doi:10.3389/fgene.2012.00246

13. Jin Y. A comprehensive survey of fitness approximation in evolutionary computation. Soft Comput. Springer-Verlag; 2005;9: 3–12. doi:10.1007/s00500-003-0328-5

14. Orgogozo V. Replaying the tape of life in the twenty-first century. Interface Focus. 2015;5: 20150057. doi:10.1098/rsfs.2015.0057

15. Niklas KJ. Adaptive walks through fitness landscapes for early vascular land plants. Am J Bot. John Wiley & Sons, Ltd; 1997;84: 16–25. doi:10.2307/2445878

16. Perttunen J, Sievänen R, Nikinmaa E. LIGNUM: a model combining the structure and the functioning of trees. Ecol Modell. Elsevier; 1998;108: 189–198. doi:10.1016/S0304-3800(98)00028-3

17. Fourcaud T, Blaise F, Lac P, Castéra P, de Reffye P. Numerical modelling of shape regulation and growth stresses in trees. Trees. Springer-Verlag; 2003;17: 31–39. doi:10.1007/s00468-002-0203-5

18. Nikinmaa E, Messier C, Sievanen R, Perttunen J, Lehtonen M. Shoot growth and crown development: effect of crown position in three-dimensional simulations. Tree Physiol. Oxford University Press; 2003;23: 129–136. doi:10.1093/treephys/23.2.129

19. Honda H. Description of the form of trees by the parameters of the tree-like body: Effects of the branching angle and the branch length on the shape of the tree-like body. J Theor Biol. Academic Press; 1971;31: 331–338. doi:10.1016/0022-5193(71)90191-3

20. Graus RR, Macintyre IG. Light control of growth form in colonial reef corals: Computer simulation. Science (80-). 1976;193: 895–897. doi:10.1126/science.193.4256.895

21. Merks RMH, Hoekstra AG, Kaandorp JA, Sloot PMA. Polyp oriented modelling of coral growth. J Theor Biol. 2004;228: 559–576. doi:10.1016/j.jtbi.2004.02.020

22. Sentoku A, Ezaki Y. Regularity in budding mode and resultant growth morphology of the azooxanthellate colonial scleractinian Tubastraea coccinea. Coral Reefs. Springer-Verlag; 2012;31: 67–74. doi:10.1007/s00338-011-0808-5

23. Ohno R, Sentoku A, Ezaki Y, Masumoto S. Modelling and Simulation of Morphogenesis in Colonial Azooxanthellate Scleractinians. Geoinformatics. 2016;27: 3–12. doi:10.6010/geoinformatics.27.1_3

24. Honda H, Fisher JB. Tree branch angle: maximizing effective leaf area. Science. American Association for the Advancement of Science; 1978;199: 888–90. doi:10.1126/science.199.4331.888

25. Honda H, Fisher JB. Ratio of tree branch lengths: The equitable distribution of leaf clusters on branches. Proc Natl Acad Sci U S A. National Academy of Sciences; 1979;76: 3875–9. doi:10.1073/PNAS.76.8.3875

26. Kanemaru N, Chiba N, Takahashi K, Saito N. Simulation of Natural Shapes of Botanical trees Based on Heliotropism. IEICE Trans Inf Syst. 1992;75: 76–85.

27. Koike F. Foliage-Crown Development and Interaction in Quercus Gilva and Q. Acuta. J Ecol. 1989;77: 92. doi:10.2307/2260919

28. Sorrensen-Cothern KA, Ford ED, Sprugel DG. A Model of Competition Incorporating Plasticity through Modular Foliage and Crown Development. Ecol Monogr. John Wiley & Sons, Ltd; 1993;63: 277–304. doi:10.2307/2937102

29. Takenaka A. A simulation model of tree architecture development based on growth response to local light environment. J Plant Res. Springer-Verlag; 1994;107: 321–330. doi:10.1007/BF02344260

30. Yoshizawa D, Yokozawa H. Trees Growth Modeling in Consideration of the Photoenvironment. J Graph Sci Japan. 2007;41: 3–9. doi:10.5989/jsgs.41.3_3

31. Onoda Y, Wright IJ, Evans JR, Hikosaka K, Kitajima K, Niinemets Ü, et al. Physiological and structural tradeoffs underlying the leaf economics spectrum. New Phytol. John Wiley & Sons, Ltd (10.1111); 2017;214: 1447–1463. doi:10.1111/nph.14496

32. Fogle CA, Nagle JL, Desai MM. Clonal interference, multiple mutations and adaptation in large asexual populations. Genetics. 2008;180: 2163–2173. doi:10.1534/genetics.108.090019

33. Keller TE, Wilke CO, Bull JJ. INTERACTIONS BETWEEN EVOLUTIONARY PROCESSES AT HIGH MUTATION RATES. Evolution (N Y). John Wiley & Sons, Ltd (10.1111); 2012;66: 2303–2314. doi:10.1111/j.1558-5646.2012.01596.x

34. Franke J, Klözer A, de Visser JAGM, Krug J. Evolutionary Accessibility of Mutational Pathways. Wilke CO, editor. PLoS Comput Biol. Public Library of Science; 2011;7: e1002134. doi:10.1371/journal.pcbi.1002134

35. Yokozawa M, Kubota Y, Hara T. Crown architecture and species coexistence in plant communities. Ann Bot. Oxford University Press; 1996;78: 437–447. doi:10.1006/anbo.1996.0140

36. Fox JF. Alternation and Coexistence of Tree Species. Am Nat. University of Chicago Press; 1977;111: 69–89. doi:10.1086/283138

37. Grubb PJ. THE MAINTENANCE OF SPECIES-RICHNESS IN PLANT COMMUNITIES: THE IMPORTANCE OF THE REGENERATION NICHE. Biol Rev. John Wiley & Sons, Ltd (10.1111); 1977;52: 107–145. doi:10.1111/j.1469-185X.1977.tb01347.x

